# Systems levels analysis of lipid metabolism in oxygen-induced retinopathy

**DOI:** 10.1101/2023.11.21.568200

**Authors:** Charandeep Singh

## Abstract

Hyperoxia induces glutamine-fueled anaplerosis in the Muller cells, endothelial cells, and retinal explants. Anaplerosis takes away glutamine from the biosynthetic pathway to the energy-producing TCA cycle. This process depletes biosynthetic precursors from newly proliferating endothelial cells. The induction of anaplerosis in the hyperoxic retina is a compensatory response, either to decreased glycolysis or decreased flux from glycolysis to the TCA cycle. We hypothesized that by providing substrates that feed into TCA, we could reverse or prevent glutamine-fueled anaplerosis, thereby abating the glutamine wastage for energy generation. Using an oxygen-induced retinopathy (OIR) mouse model, we first compared the difference in fatty acid metabolism between OIR-resistant BALB/cByJ and OIR susceptible C57BL/6J strains to understand if these strains exhibit metabolic difference that protects BALB/cByJ from the hyperoxic conditions and prevents their vasculature in oxygen-induced retinopathy model. Based on our findings from the metabolic comparison between two mouse strains, we hypothesized that the medium-chain fatty acid, octanoate, can feed into the TCA and serve as an alternative energy source in response to hyperoxia. Our systems levels analysis of OIR model shows that the medium chain fatty acid can serve as an alternative source to feed TCA. We here, for the first time, demonstrate that the retina can use medium-chain fatty acid octanoate to replenish TCA in normoxic and at a higher rate in hyperoxic conditions.

## Introduction

Premature infants are born with underdeveloped lungs, which necessitates the use of supplemental oxygen to prevent mortality in these infants[1–6]. Supplemental oxygen prevents mortality but also leads to the development of retinal vascular abnormality-retinopathy of prematurity (ROP). Supplemental oxygen (hyperoxia) downregulates angiogenesis in phase I and causes vaso-obliteration, a precursor to phase II[6]. Every year, ROP claims the eyesight of nearly 30,000-40,000 prematurely born infants[7]. Multiple biochemical pathways have been implicated in the ROP and mouse oxygen-induced retinopathy (OIR) model, including serine-one carbon, glutamine metabolism, amino acid metabolism, polyamine pathway, and fatty acid metabolism[6]. These metabolic changes in the oxygen-induced retinopathy model and the ROP indicate a biosynthetic deficit. Proliferating endothelial cells have high biosynthetic demand to proliferate and migrate to develop new blood vessels. Inhibition of endothelial cell (EC)-specific glutaminase (GLS) has been demonstrated to decrease proliferation and induce senescence in ECs[8, 9]. In addition, ablation of glutamine synthetase in Muller cells has been shown to cause EC proliferation defects[10]. This shows that glutamine production and utilization by the retina need to be balanced during angiogenesis. Similarly, loss of serine transporter from the retina demonstrates aberrant angiogenesis in the retina[11]. In addition, polyamine pathway metabolites have also been shown to cause neovascular defects in the retinal and retinal cell types[12, 13]. These findings demonstrate that biosynthetic pathways are essential during angiogenesis. Glycolysis and OXPHOS provide ATP necessary for cellular function; however, biosynthetic pathways like serine-one carbon, glutamine and polyamine pathways are necessary for the cells to divide. At the cellular level, serine-one carbon metabolites are essential for glutathione production, NADPH synthesis, and Heme production pathways[2]. Serine-one carbon is the only source of methylation in the early developmental stages[14]. At the same time, glutamine is an essential nutrient for cellular proliferation[10]. Glutamine and downstream urea cycle metabolites arginine and ornithine also serve as precursors for polyamine pathway metabolites[15]. Glutamine is present in millimolar ranges, both in humans and mice, and because of its high abundance, it also serves as an alternative source of energy. However, glutamine filling into TCA is bad for proliferating tissue as this lowers the amount available for biosynthetic reactions and produces ammonium, a toxic end product. Carbon sources that can produce energy via the TCA cycle, other than glycolysis and glutaminolysis, are fatty acids and branched amino acids. Newborns rely on fatty acids as their main energy source, as fats account for 40-50% of calories in the mother’s milk[16]. Free fatty acid also contributes to the majority of energy generation during fetal development[16]. During development, the fetus is supplied with free fatty acids from the maternal circulation. Bitman et. al. have demonstrated that the fatty acid composition of the mother’s milk in premature and full-term infants differs in medium chain fatty acids (MCFAs)[17]. The highest amounts of MCFAs were observed in the mother’s milk of very preterm-born infants[16]. MCFAs are a very good energy source as they don’t need a carnitine-based transport to be oxidized, unlike required for long and very long chain fatty acids in many tissues, except for a few tissue types[18]. In addition, MCFAs are broken down by the action of lingual, milk, and gastric lipases and don’t need to rely on pancreatic lipase, which is low during the early stages of life[16]. This makes MCFAs a preferred energy source during the early stages of life.

The liver plays an important role in managing fatty acids in the body, both the production and the transport to different organs. We have demonstrated that the liver-based HIF1α is important to produce serine-one carbon and remotely provide biosynthetic products to the retina[2]. In addition, signaling molecules like FGF21, a hepatokine, have been demonstrated to regulate physiological and pathological angiogenesis[19]. These findings demonstrate a system’s level of metabolic and biochemical signaling in retinal angiogenesis during the early stages of life. Unlike genetic disorders, where the gene product of one or more pathways contributes to pathology, the ROP develops solely due to a change in oxygenation level in the environment to which the whole infant is exposed. Although the whole body is exposed to hyperoxia, most of the studies are focused locally on the retina. We used oxygen-induced retinopathy model to investigate systems-level liver-retina metabolic exchanges in response to hyperoxia. This model recapitulates vaso-obliteration and neovascularization as seen in human disease ROP. Ritter et al. have demonstrated that the BALB/cByJ strain is resistant to the OIR[20]. In this publication, we performed a multi-omics interorgan comparison of the OIR-resistant and OIR-susceptible strains of mice to find the genetic basis of protection.

## Results

### C57BL/6J mouse strain is susceptible, whereas BALB/cByJ is resistant to oxygen-induced retinopathy

Ritter et al. have previously published that the BALB/cByJ mice are resistant to OIR and have normal vasculature when exposed to the OIR model[20]. We repeated the experiment and investigated the vasculature of C57BL/6J and BALB/cByJ mice exposed to oxygen-induced retinopathy models in phase 1 and phase 2. We used oxygen induced retinopathy model as in Smith et. al.[21]. Mice were maintained in room air from postnatal day 0 (P0) to postnatal day 7(P7). On P7, mice were moved to 75% oxygen until P12. On P12, mice were sacrificed to investigate phase 1 phenotype. Some mice were again brought back to room air and maintained in room air from P12-P17, which concluded the model on P17 and phase 2.

Mouse strain C57BL/6J showed a clear avascular region in the middle of the retina, whereas this region was almost negligible in the BALB/cByJ strain in phase 1 of OIR (figure 1 a,b,d, and e). C57BL/6J further developed neovascular tufts and tortuosity in the vasculature in phase 2 of OIR on P17, as seen in figure 1c. In contrast, the BALB/cByJ retina continued to develop a healthy vasculature, as seen in figure 1f on P17. This difference in susceptibility to OIR demonstrates the genetic basis of protection in BALB/cByJ mice strains.

**Figure 1.**
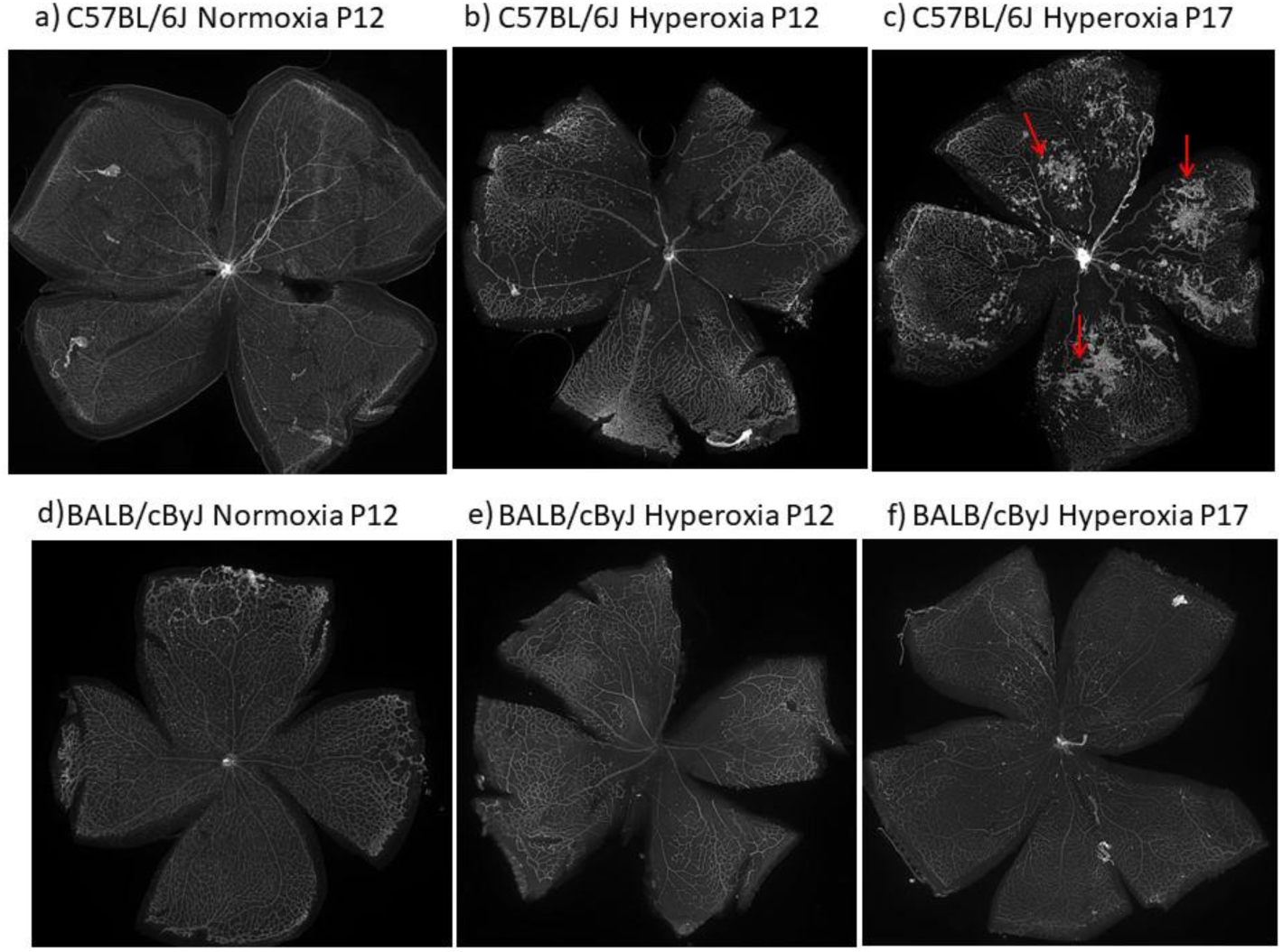
C57BL/6J is susceptible to oxygen induced retinopathy whereas BALB/cByJ is resistant to oxygen induced retinopathy. a,d, normoxic condition P12; b,e, mice exposed to hyperoxia from P7-P12; c,f, mice exposed to hyperoxia from P7-P12 and then room air until P17.

### RNA sequencing data indicates the downregulation of biosynthetic pathways in OIR susceptible strains-C57BL/6J as compared to the OIR resistant strain-BALB/cByJ, and the upregulation of medium-chain fatty acid utilization pathway in the OIR resistant strain-BALB/cByJ

To understand the genetic basis of protection, we looked at the transcriptional regulation of metabolic pathways that protect against OIR by comparing the retinal transcriptome of an OIR-susceptible strain to an OIR-resistant strain of mice. We performed RNA sequencing on the retinas isolated from these two mouse strains in hyperoxic conditions. Mice were incubated in hyperoxia following the OIR model[21], retinas were dissected on P12, and RNA sequencing was performed. We first investigated the biosynthetic pathway, which we have demonstrated to differ between diseased and control retinas during phase 1 in our previous publications[2, 6, 13, 22]. Interestingly, we found differences in genes involved in the serine/one-carbon uptake and utilization, glycolytic regulator PDK which controls the entry of glucose into TCA, and polyamine pathway (figure 2 b, c, and d). Since hyperoxia leads to glutamine-fueled anaplerosis, we looked at the fatty acid breakdown pathways that can feed into the TCA cycle in the absence of flux from glycolysis or glutamine breakdown. There are only a few alternative pathways that can feed into TCA. These pathways include fatty acid oxidation, branched-chain amino acids, glutamine, and proline breakdown. Of all these pathways, medium-chain fatty acids have been shown to differ in the milk of premature infant mothers, which implies their importance in ROP[17]. We looked at all the acyl-coA dehydrogenase, which is the first step to break down each fatty acid type (short, medium, long and very long chain), and we only found statistically significant changes in Acadm, which breaks down medium-chain fatty acids. We found higher expression of these genes in the OIR-resistant strain than that of the susceptible strain; this included Acadm, Ech1 and Acaa2 (figure 2a). This pathway breaks down medium-chain fatty into their corresponding Acyl Coa derivative, which can then feed into TCA and provide energy when the entry of carbon from other sources into TCA is limited. In addition, we also found upregulation of methylmalonyl-CoA mutase, which is required to facilitate the entry of odd-chain fatty acids and branched-chain amino acids into the TCA.

**Figure 2.**
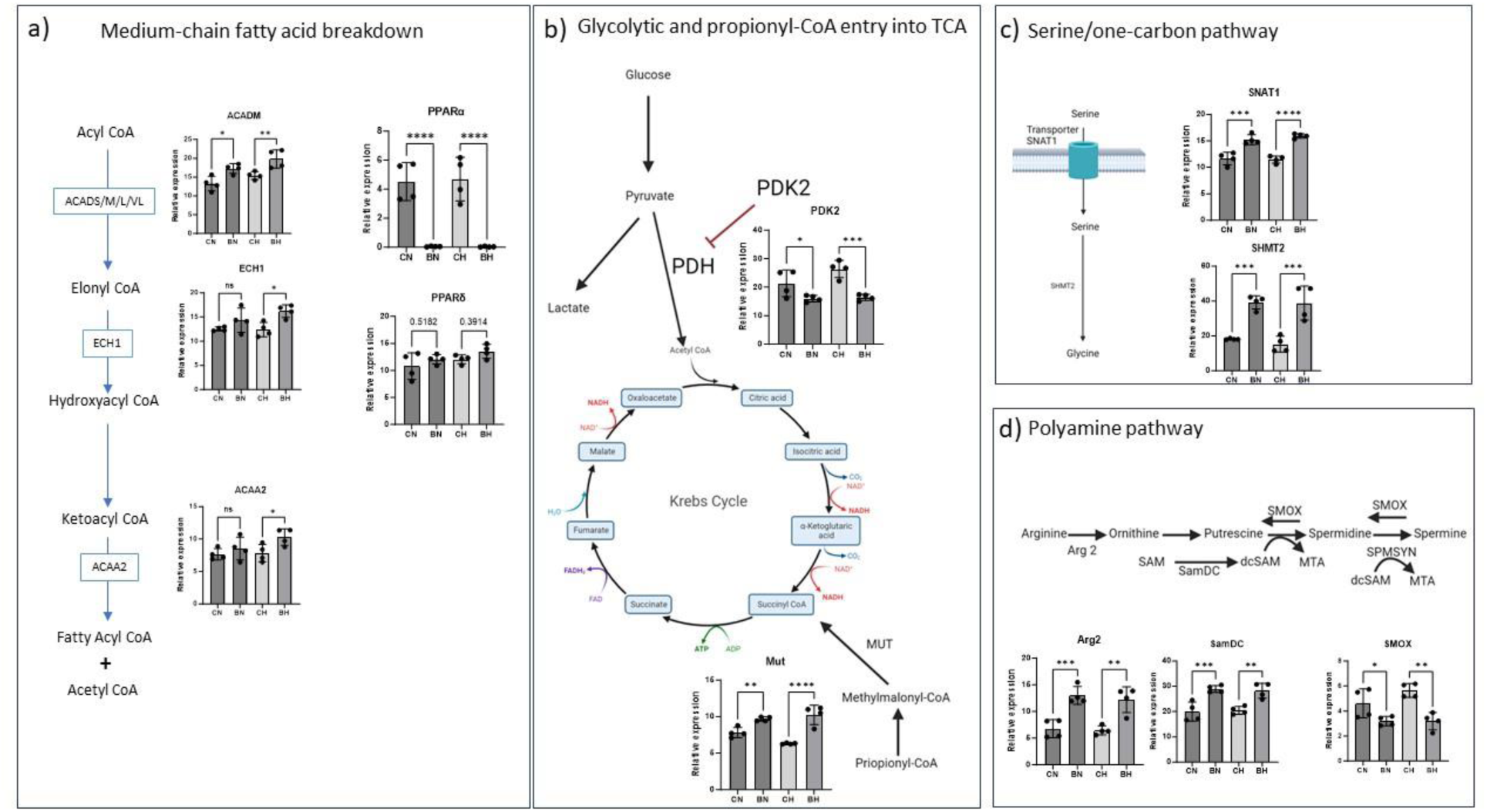
Retinal gene expression changes in P12 mice a) Changes in medium chain fatty acid utilization enzymes ACADM, ECH1, and ACAA2. Retinal PPARα negatively correlates with the expression of medium-chain fatty acid utilization genes. PPARδ changes were not significant between two mouse strains compared. b) Glycolytic inhibitor PDK2 which inhibits PDH and inhibits flux from glycolysis entering TCA showed statistically significant changes between the two mouse strains. MUT is an additional entry point of propionyl-CoA and branch-chain amino acids. MUT showed statistically significant changes between the two strains. P-values were calculated using unpaired one-way ANOVA with Bonferroni corrections applied for multiple comparison, * p-value <=0.05, ** p-value <=0.01, *** p-value <= 0.001 and **** p-value <=0.0001. Legends: CN, C57BL/6J normoxia; BN, BALB/cByJ normoxia; CH, C57BL/6J hyperoxia; BH,BALB/cByJ hyperoxia.

### Secretome data indicates differential changes in growth factors in circulation between two mouse strains exposed to hyperoxia

Since growth factors like FGF[19, 23] and IGF[24–26] have been shown to correlate with developmental angiogenesis and have been shown to correlate with phase 1 pathogenesis in oxygen-induced retinopathy, we investigated these factors using the proteomics platform SOMAScan. This platform utilizes aptamer-based technology, which very specifically binds protein targets with the right structure, therefore avoiding non-specific binding like seen in other proteomics techniques. In addition, this technique has a better dynamic range as compared to other proteomics techniques. This dynamic range is specifically beneficial for plasma samples in which albumin and other high abundance proteins hinder the measurement of low abundance targets.

IGF1 and its binding partner IGF ALS both correlated with the pathology. These proteins were downregulated in C57BL/6J mice exposed to hyperoxia. We also investigated IGFBPs and discovered that the IGFBP-5 correlated with the IFG1 and IGF ALS. Additionally, the expression of IGFP-2 expression increased in response to hyperoxia in both the C57BL/6J and BALB/cByJ strains. IGFBP-1 and IGFBP-3 were moderately altered in both the strains. These findings are presented in figure 3.

**Figure 3.**
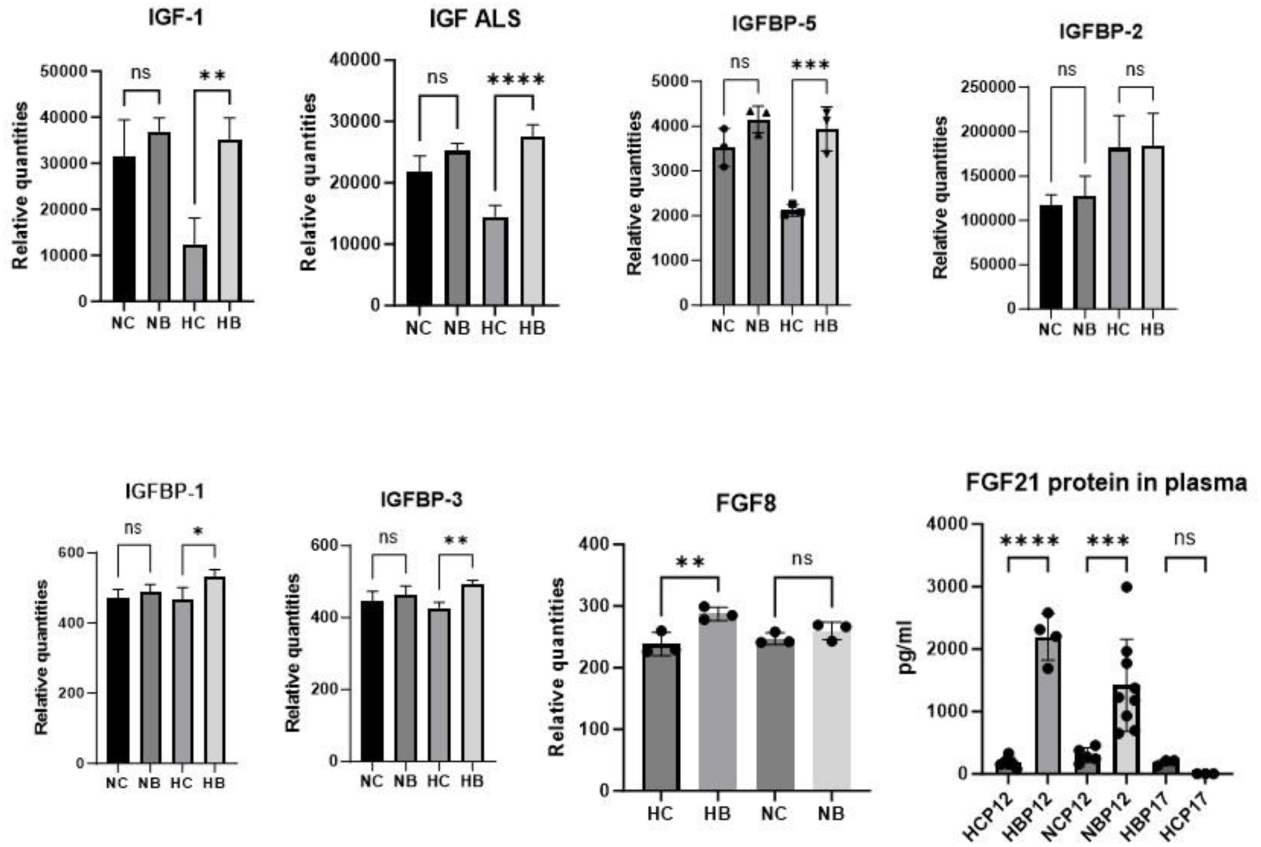
Circulating IGFs, IGFBPs, and FGFs show statistically significant difference between the two strains, and these changes correlate with protection against OIR. p-values were calculated using unpaired one-way ANOVA with Bonferroni corrections applied for multiple comparison, * p-value <=0.05, ** p-value <=0.01, *** p-value <= 0.001 and **** p-value <=0.0001. Legends: NC, P12 C57BL/6J normoxia; NB, P12 BALB/cByJ normoxia; HC, P12 C57BL/6J hyperoxia; HB, P12 BALB/cByJ hyperoxia; HCP12, P12 C57BL/6J hyperoxia; HBP12, P12 BALB/cByJ hyperoxia; NCP12, P12 C57BL/6J normoxia; NBP12, P12 BALB/cByJ normoxia; HBP17, P17 BALB/cByJ hyperoxia; HCP17, P17 C57BL/6J hyperoxia.

Since FGFs have previously been shown to contribute to oxygen-induced retinopathy, we measured FGF1/bECGF, FGF2/bFGF, FGF3, FGF4, FGF5, FGF6, FGF7, FGF8, FGF8A, FGF8B, FGF8F, FGF9, FGF10, FGF12, FGF16, FGF17, FGF18, FGF19, FGF20, FGF22, FGF23, FGFP1 and FGFP3 using SomaScan. Among these FGF and FGF binding proteins, only FGF8 showed statistically significant difference between conditions compared. Since FGF21 has been shown to promote vascularization in phase 1 ROP via lipid oxidation, we investigated the protein levels of FGF21 in both the mice strains with the help of an ELISA kit because FGF21 is not among the SOMAscan targets. These findings are also presented in figure 3. FGF21 showed statistically significant drastic differences between C57BL/6J and BALB/cByJ mice exposed to hyperoxia.

### Genes required to produce ketone body and β-oxidation products in the liver were upregulated in the BALB/cByJ compared to C57BL/6J

### Liver

Since medium chain fatty acids are known to be cleaved in the liver as well. We investigated genes which uptake fatty acids from circulation. Fatty acids are taken up by the liver specific FABPs, and then broken down into ketone bodies. These ketone bodies can serve as a source of energy for peripheral tissues. This process also releases non-esterified fatty acids which can leak through the liver into the circulation and serve as an additional source of energy. Liver specific PPARα and PPARβ/δ showed higher expression in BALB/cByJ exposed to hyperoxia as compared to C57BL/6J. In addition, targets downstream of PPARα, involved in beta oxidation, demonstrated higher expression in the BALB/cByJ exposed to hyperoxia in comparison to C57BL/6J. All the expression data is provided in figure 4.

**Figure 4.**
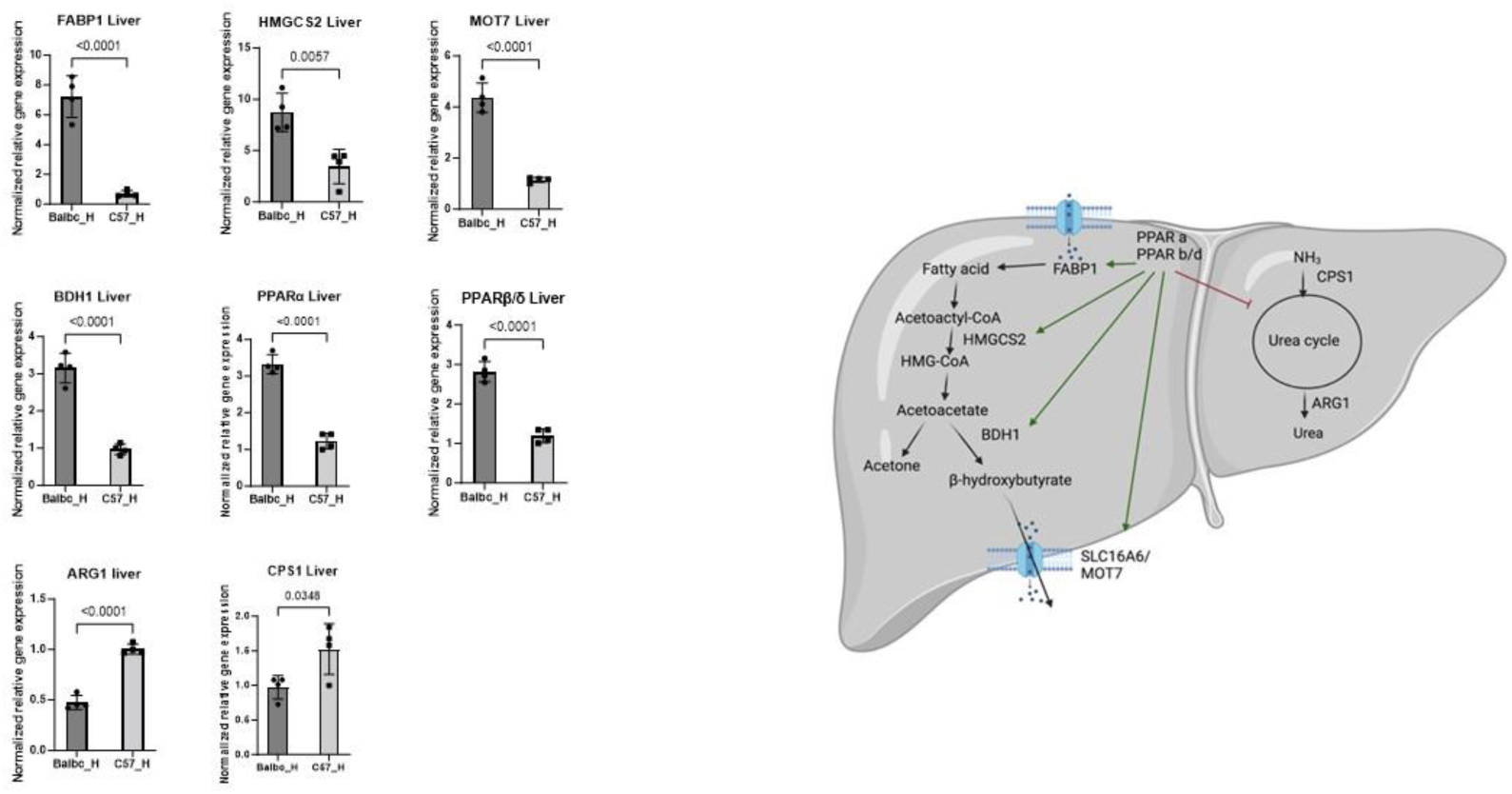
Liver gene expression changes in fatty acid breakdown pathways in P12 old mice shows higher expression of enzymes in fatty acid oxidation and transport of ketone bodies in OIR-resistant strain of mice, whereas urea cycle genes demonstrate higher expression in OIR-susceptible mice indicative of difference in substrate utilization for energy. p-values were calculated using t-test. Legends: Balbc_H, P12 BALB/cByJ hyperoxia; C57_H, P12 C57BL/6J hyperoxia.

In addition, urea cycle genes, ARG1 and CPS1, which are inhibited by PPARα showed lower expression in BALB/cByJ as compared to C57BL/6J (figure 4), implying the lower utilization of nitrogenous compounds for energy production. Nitrogenous compounds, like glutamine, release ammonium when deaminated or deamidated to produce energy via TCA cycle, which necessitates the use of liver-based urea cycle to remove toxic ammonium.

We additionally performed an RNA sequencing analysis on the RNA extracted from the liver samples taken from two strains exposed to normoxic or hyperoxic conditions, to look at additional genes which were not evaluated in the qPCR analysis. Since, hepatic FGF21 has been shown to control lipid metabolism in the body via liver-brain axis[27], we first looked at hepatic FGF21 expression levels in the hepatic RNA sequencing data to determine if FGF21 levels in the plasma correlate with the hepatic FGF21 in the OIR model at P12. Liver and milk are the two main sources of FGF21 in the early stages of life[28]. Since the source of FGF21 protein in the plasma in OIR is not clear, we performed correlation analysis of hepatic FGF21 transcript and plasma FGF21 proteins levels. We found a very strong correlation between hepatic FGF21 transcript levels measured using RNA sequencing and plasma levels of FGF21 measured with an ELISA (figure 5). We additionally looked at some of the known inducers of FGF21 in the liver that have been previously reported. PPARα is a known master regulator of FGF21[29]. High fat diet, ketogenic diet, and fasting are the main activators of PPARα dependent FGF21 induction[29]. CREBH shows some cross talk with the PPARα during overnutrition and excess lipid availability[30]. Transcript levels of FGF21 in the liver strongly correlated with the transcript levels of PPARα and CREBH in our RNA seq data (figure 5). Some of the other known transcriptional inducers of FGF21 in the liver are ChREBP, ATF4, Nurp1, and XBP1[30]. ChREBP is a carbohydrate sensitive transcription factor, and it induces hepatic FGF21 in response to glucose, fructose, and ethanol. Whereas ATF4 and Nurp1 activate FGF21 in response to changes in amino acid concentrations in the liver[31–33]. Some studies have suggested that the ER stress involves XBP1 to induce FGF21, however, recent findings demonstrate XBP1 is dispensable and may not be the direct regulator of FGF21 in the mouse liver[32]. In our correlation analysis, ChREBP, ATF4, Nurp1, and XBP1 transcripts negatively correlated with the FGF21 levels (figure 5). Correlation analysis indicates that the PPARα and CREBH contribute to the FGF21 induction in OIR context.

**Figure 5.**
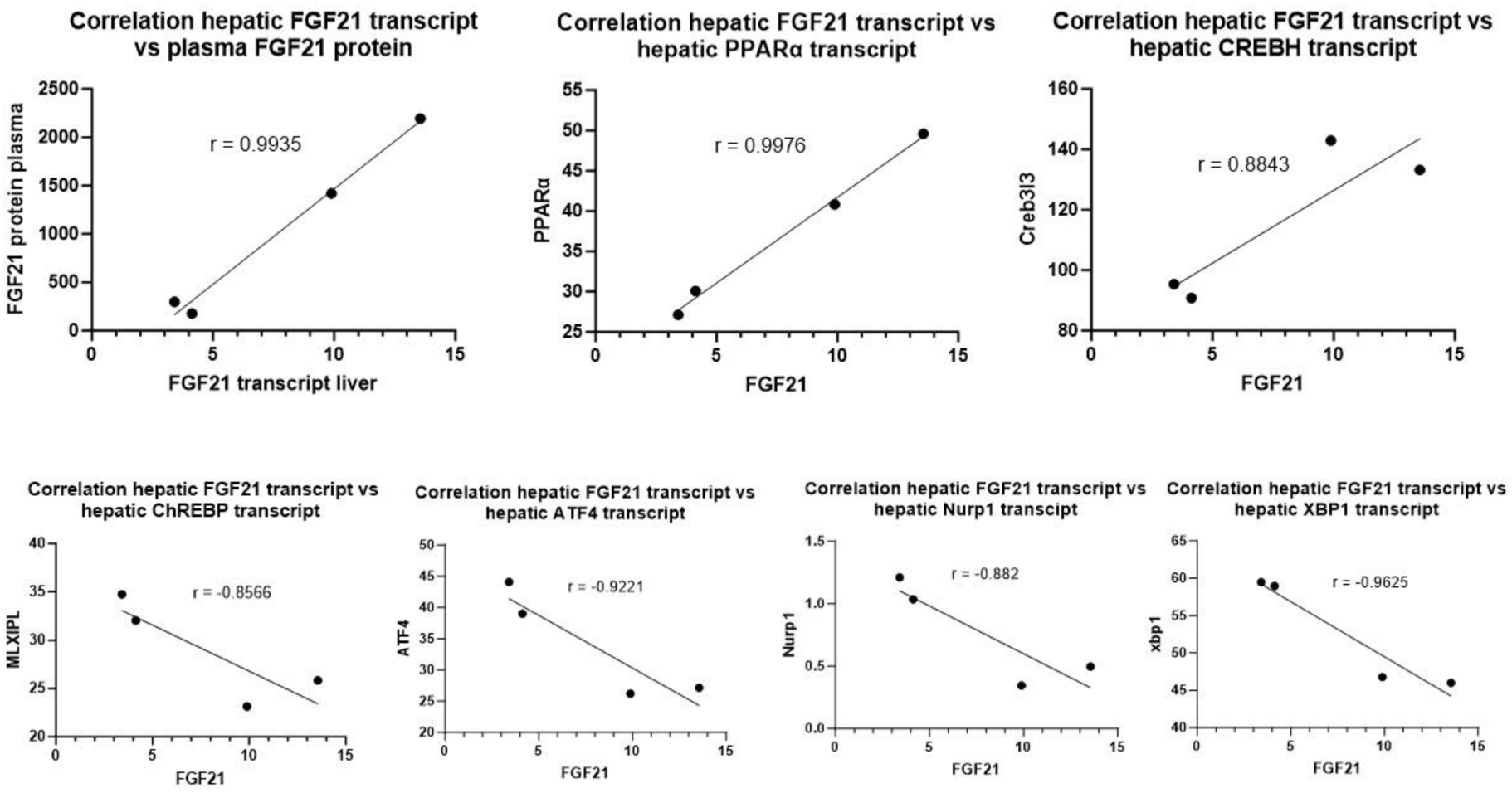
Liver based FGF21 is secreted into the plasma in P12 old mice, and its induction involves hepatic PPARα and CREBH. Hepatic FGF21 transcript levels strongly correlate with FGF21 protein levels in plasma. Hepatic FGF21 transcript levels also strongly correlate with transcript levels of PPARα and CREBH. Hepatic transcript levels of ChREBP, ATF4, Nurp1, and XBP1, negatively correlated with the transcript levels of FGF21 in the liver.

### Stable isotope labeling of retinal explant indicates octanoate can replenish TCA cycle in response to hyperoxia

Based on our findings from the RNA sequencing data and known differences in MCFA levels in the milk of preterm vs. full-term infants, we hypothesized that retina may be able to utilize medium-chain fatty acids to fill its TCA cycle in hyperoxic conditions. We incubated retinal explants from C57BL/6J mice in media containing [^13^C_8_] labeled octanoate. Retinal explants were able to use octanoate as a substrate to feed into their TCA cycle and in addition, this process was accelerated in hyperoxic conditions. This implies that medium-chain fatty acid octanoate can fuel the TCA cycle to replenish the glutamate pool. Total quantities of citrate and its M6 form were higher in the hyperoxic retinal explants as compared to normoxic explants. In addition, both total glutamate and M5 form of glutamate were higher in hyperoxic retinal explants as compared to normoxic explants (figure 6), indicative of higher conversion of [^13^C_8_] octanoate into citrate and subsequently into glutamate.

**Figure 6.**
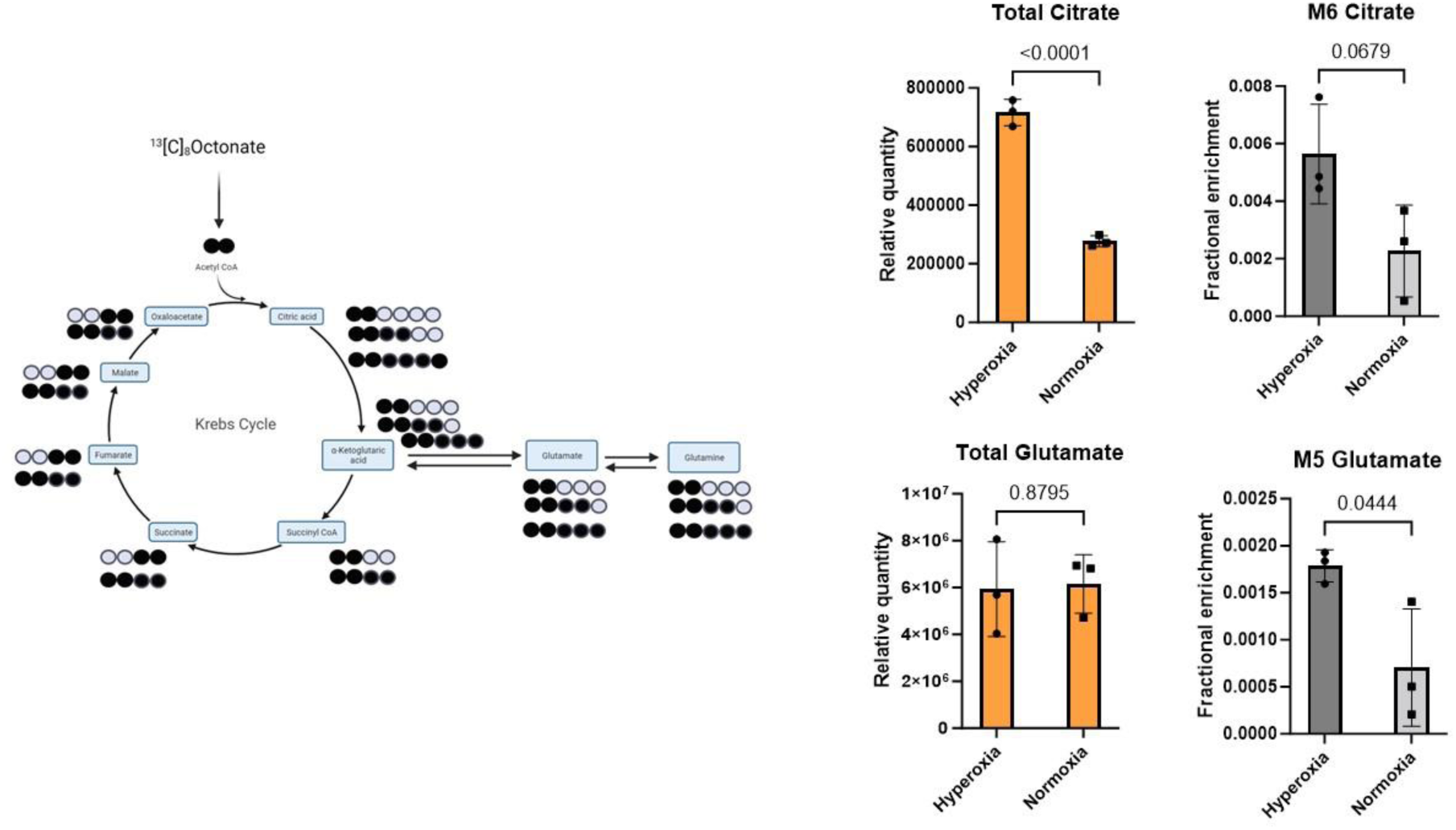
Stable isotope labeling of C57BL/6J retinal explants with [^13^C_8_] octanoate indicates retina can utilize medium chain fatty acid to replenish its TCA cycle and normalize its total glutamate levels in hyperoxic conditions. Total citrate and total glutamate are the sum of all MIDs. M6 citrate and M5 glutamate are fully labeled forms of citrate and glutamate, respectively. p-values were calculated using t-test.

## Methods

### OIR model and tissue collection

All the animals used in this work were used under Massachusetts General Hospital Institutional Animal Care and Use Committee approved protocol. The OIR protocol used was as described in the publication by Smith et. al.[21]. Briefly, mice were maintained in room air from P0-P7, on P7 mice were exposed to hyperoxia 75% oxygen until P12 which caused vaso-obliteration Phase I of OIR. P12 hyperoxic treated mice, when maintained in room air on P17 demonstrated retinal neovascularization. Control mice were kept in room air until P12 or P17. All the mice were anesthetized with isoflurane and euthanized by cervical dislocation before the removal of tissues. Blood was collected from retro-orbital route into heparin tubes and spun down at 1000x g for 20-30 min at 4 °C to separate plasma. Plasma was flash-frozen in liquid nitrogen and kept at –80 °C until use. Livers were removed from euthanized mice using forceps and immediately frozen in liquid nitrogen. Livers were ground in a mortar pestle in liquid nitrogen, and then powdered livers were stored at –80 °C for later use.

### Retina staining with Isolectin

Retinal flat mounts were prepared and stained as described in Jidigam et. al.[34]. Isolectin GS-IB4 from *Griffonia simplicifolia*, Alexa Flourtm 568 conjugate (Invitrogen) was used to label blood vessels. Images were acquired on Nikon 90i upright epifluorescence microscope at MGH microscopy core facility.

### RNA sequencing

Total RNA was extracted using TRIzol reagent (Invitrogen) as per instructions provided with the reagent. Following extraction, RNA was cleared on DNA using DNA-free DNase Removal kit (Invitrogen). The sample quality was determined on an Agilent 5400 fragment analyzer instrument. The library was prepared with the help of a NEBNext Ultra II library prep kit for Illumina. The samples were sequenced on NovaSeq 6000 with the help of NovaSeq 6000 S4 Illumina kit. Fastq formatted files were processed to remove reads containing adapters, lower quality reads and poly-N reads. Reference genome index was prepared using Hisat2 v2.0.5[35]. *Mus Musculus* reference genome GRCm38/mm10 was used as reference. Quantification of gene expression was performed with the help of featureCounts v1.5.0-p3[36]. FPKM was calculated for each gene.

### SOMAscan proteomics

Blood samples were collected via retroorbital route into heparin tubes. Samples were spun at 1000 x g at 4 °C for 20-30 min and flash frozen in liquid nitrogen. Samples were later thawed and subjected to measurement on SOMAscan7K v4.1 proteomics platform as described previously as in Haslam et. al.[37].

### Metabolite labeling and measurement

Eyes were enucleated from P12 old mice after euthanasia. Eyes were maintained in Astrocyte media (A1261301) on an ice pack during dissection. After dissection, retinas were removed from the media, washed with normal saline and then incubated in media containing [^13^C_8_] octanoate. To prepare this custom stable isotope media, first BSA bound [^13^C_8_] octanoate was prepared by dissolving 16 µl of octanoate in 1 ml of ethanol and then adding 200 µl of above in 2.5 ml of fatty acid free BSA solution (1.133 mg/ml prepared in 150 mM NaCl aqueous solution) made to 10 ml total volume with 150 mM NaCl. The BSA bound [^13^C_8_] octanoate was added in 1:5 dilution with minimal media (DMEM catalogue number A14430-01) containing 5 mM glucose, 2.5 mM glutamine and 1% PenStrep (10000U/ml stock solution, Invitrogen). Two retinas were incubated per well in 500 µl of media volume for 24 h. One plate was exposed to hyperoxia 75% oxygen and other plate was exposed to normoxia 21%, in 5% CO_2_ incubator at 37°C.

Metabolites were extracted by adding 1ml of 80% methanol per tube. Tubes were centrifuged at 15000x g for 5 min at 4°C, and the supernatant was dried for metabolite measurement. GCMS measurements were performed on 8890B GC coupled to EI/CI 5977 MSD mass spectrometer using an electron impact ionization extractor ion source (Agilent) as previously described[13].

### FGF21 ELISA

FGF21 ELISA was performed following the manufacturer’s guidelines (FGF21 ELISA kit, Millipore Sigma catalogue number-EZRMFGF21-26K).

### Correlation analysis

Two targets, protein vs. transcript or transcript vs. transcript, were plotted and pearson correlation coefficient was calculated. Statistical analysis were performed using GraphPad Prism 10 version 10.1.0.

### qPCR experiment

cDNA was prepared with the help of iScript gDNA clear cDNA synthesis kit (Biorad). cDNA was added to iTaq Universal SYBR Green Supermix and gene specific primers. PCR settings used were as previously described in Singh et. al.[6]

Sequences of the PCR primers used

PPARα fwd 5’ ACGATGCTGTCCTCCTTGAT 3’

PPARα rev 5’ GATGTCACAGAACGGCTTCC 3’

PPAR β/δb/d fwd 5’ TCAAGTTCAATGCGCTGGAG 3’

PPAR β/δ rev 5’ TGTCCTGGATGGCTTCTACC 3’

CPS1 fwd 5’ GTGAAGGTCTTGGGCACATC 3’

CPS1 rev 5’ TTCCACTGCAAAACTGGGTG 3’

ARG1 fwd 5’ AGAGATTATCGGAGCGCCTT 3’

ARG1 rev 5’ AGTTTTTCCAGCAGACCAGC 3’

MOT7 fwd 5’ATACAGCCTCCTCTTCGTGG 3’

MOT7 rev 5’ CAGAGTGACTGCTTTCGGTG 3’

HMCS2 fwd 5’ GCATAGATACCACCAACGCC 3’

HMCS2 rev 5’ TCGGGTAGACTGCAATGTCA 3’

FABP1 fwd 5’ CTTCTCCGGCAAGTACCAAT 3’

FABP1 rev 5’ TCCCTTTCTGGATGAGGTCC 3’

BDH1 fwd 5’ GGATTTGGGTTCTCACTGGC 3’

BDH1 rev 5’ CCACCTCTTCACTGTTGCAG 3’

GAPDH fwd 5’ CAACGACCCCTTCATTGACC 3’

GAPDH rev 5’ TCCTGGAAGATGGTGATGGG 3’

## Discussion

In this study we demonstrate in multiple ways that OIR is not limited to the retina, but rather a systemic disease. In this systems levels comparison of an OIR-resistant and susceptible strain, we first confirmed our previous findings and demonstrated that OIR is a biosynthetic deficit. We saw higher expression of serine pathway genes, polyamine biosynthesis genes in retina of OIR-resistant strain as compared to OIR-susceptible strain of mice. In addition, we also observed lower expression of PDK2, which inhibits PDH and blocks entry of glycolytic carbon into TCA cycle, in the retina of OIR-resistant strains as compared to OIR-susceptible strain. In addition, we also observed lower expression of polyamine breakdown enzyme SMOX in OIR resistant mice in comparison of OIR-susceptible mice. This implies that the OIR resistant mice can preserve its biosynthetic molecules for proliferation of endothelial cells in developing vasculature in the retina. This may be in response to availability of alternative source of energy other than using these biosynthetic pathway metabolites for energy production via TCA cycle. In our previous publication we have demonstrated that hyperoxia induces glutamine-fueled anaplerosis which depletes glutamine from biosynthetic pathways and reroutes it towards its oxidation via TCA cycle[22]. Some of the alternative sources of TCA filling are fatty acids and branched chain amino acids. In this publication, we focused on systems levels analysis of utilization and transcriptional regulation of fatty acids. Fatty acids are highly abundant in milk and 40-50% of calories a newborn consumes comes from the fatty acids present in mother’s milk[16]. Medium chain fatty acid quantity in mother’s milk differs between a preterm and full-term infant implying they might have an important role in OIR and ROP. In addition, medium chain fatty acid metabolism by astrocytes in the brain has been shown to promote GABA synthesis in neurons by modulating glutamine supply[38]. Our analysis shows that the retina of OIR resistant strain has higher expression of MCFA breakdown enzymes, which breakdown MCFAs via retinal β-oxidation. In addition, [^13^C_8_] octanoate labeling of retinal explant demonstrates that the retina can use octanoate to replenish its TCA cycle. These findings suggest that MCFAs may cure biosynthetic deficit seen in phase 1 OIR. Triheptanoin, a triglyceride of 7 carbon containing medium chain fatty acid, has successfully been applied for many applications to replenish TCA cycle metabolite and cure energy crisis [39, 40]. Our data indicates that this approach may also help to prevent energy crisis seen in OIR.

One of the known transcriptional factors that induce fatty acid oxidation is PPARα[41]. Interestingly expression of PPARα negatively correlated with expression MCFA utilization enzymes in the retina in our study. FGF21, which is another well-known inducer of β-oxidation[42, 43], has been demonstrated to induce retinal angiogenesis in hyperglycemic model of OIR[19]. Interestingly, we observed higher expression of PPARα in the liver of P12 old BALB/cByJ mice, which correlates with the fatty acid import, export, and oxidation enzymes in the liver of BALB/cByJ mice; PPARα expression in the liver correlates positively with the β-oxidation enzymes in the retina too. It is possible that during the early stages of life, the liver based PPARα controls MCFA oxidation in the retina by inducing FGF21 in the liver and secreting it into the blood. Our correlation analysis demonstrated a very strong correlation between hepatic FGF21 transcript levels and plasma FGF21 protein levels, indicating FGF21 necessary for protection is secreted from the liver. Our findings that the hepatic PPARα is higher in the BALB/cByJ mice, and it is induced in response to hyperoxia, and hepatic FGF21 transcript levels and plasma FGF21 levels strongly correlate, demonstrates that hepatic PPARα may be the stimulus for FGF21 secretion in OIR context. Our finding that FGF21 only peaks up during the early stages of life, i.e higher in P12 mice as compared to that in P17 mice, indicates that the liver derived FGF21 is a developmental hepatokine required during early development of retina. FGF21 has been shown to be present in mother’s milk. The concentration of FGF21 in plasma and the milk in mouse and rat are almost comparable. In comparison, human milk has approximately half the quantity of FGF21 in milk compared to plasma. Our data demonstrates that the FGF21 protein levels in the plasma and FGF21 transcript levels (data not shown) in the retina showed no correlation. However, there was a very strong correlation between hepatic FGF21 expression in the liver and plasma FGF21 protein levels, indicative of hepatic origin of FGF21 in early stages of life. Our correlation analysis further indicates CREBH and PPARα based induction of hepatic FGF21 expression and induces secretion into the plasma. One of the known endogenous activators of PPARα is 7(S)-Hydroxydocosahexaenoic acid which is a derivative of DHA[44]. DHA levels have been demonstrated to negatively correlate with severity of ROP [45]. Pharmaceutical treatment with PPARα agonist fibrates or hepatic overexpression of PPARα using genetic tools like AAVs may prevent OIR by modulating downstream FGF21 and retinal β-oxidation. Fenofibrate has been demonstrated to protect against DR and NV in the phase II of the OIR rat model[46]. Since PPARα induces ketone body formation in liver, and our data indicates upregulation of ketone body production and secretion genes, a ketone body transport from liver to the retina may be an alternative mechanism of protection in OIR. FGF21 is not the only molecule that changes with age. There are other examples like OAT, which serves as a bidirectional enzyme in neonates and unidirectional in later stages of life[47]. The reason for this reversibility in neonatal OAT is low levels of arginine in the mother’s milk, which makes it impossible to support polyamine pathway flux for proliferative processes. Given that the mother milk contains MCFA, FGF21, and their trend changes right after birth, it warrants looking into the role of maternal nutrition in the OIR and ROP.

## Acknowledgements

This project is supported by a National Institute of Health/National Eye Institute R21 grant to CS (Grant # R21EY033046). CS performed all the experiments, analyzed the data and wrote the manuscript.

I want to thank my mentor Dr. Russell Goodman who helped immensely in making sure that I ran this project independently and yet supported in each step of this study. I want to thank Dr. Nirajan Shrestha for proofreading the manuscript and valuable suggestions. I want to additionally thank Dr. Fumito Ichinose and Dr. Eizo Marutani who allowed me to use their hyperoxia chamber for initial experiments until I obtained my own instrument at MGH. I want to thank Dr. Towia Libermann and Dr. Simon Dillon BIDMC with their help in SOMAscan measurements.

Figures were prepared using BioRender.com. Data was plotted using Graphpad prism. Microscopic Imaging was performed in the Microscopy Core of the Program in Membrane Biology, which is partially supported by a Centre for the Study of Inflammatory Bowel Disease Grant 5P30DK043351 and a Boston Area Diabetes and Endocrinology Research Center (BADERC) Award 1P30DK135043. If confocal systems were used, please add the following sentences. The AX confocal system is supported by NIH grant S10 OD032211-01 and the Zeiss confocal system by grant 1S10OD021577-01.

